# Organization in complex brain networks: energy distributions and phase shift

**DOI:** 10.1101/522797

**Authors:** Saurabh Kumar Sharma, Soibam Shyamchand Singh, Dineshchandra Haobijam, Md. Zubbair Malik, R.K. Brojen Singh

## Abstract

The Hamiltonian function of a network, derived from the intrinsic distributions of nodes and edges, magnified by resolution parameter has information on the distribution of energy in the network. In brain networks, the Hamiltonian function follows hierarchical features reflecting a power-law behavior which can be a signature of self-organization. Further, the transition of three distinct phases driven by resolution parameter is observed which could correspond to various important brain states. This resolution parameter could thus reflect a key parameter that controls and balances the energy distribution in the brain network.

---

Brain is a complex universe of neurons in which its functional and structural network properties have diverse features, such as, small worldness (minimizing wiring costs) [1], significant emergence of modules (local signal processing and propagation) [2], and a distribution of rich club of key hubs (crucial for global signal integration) [3]. These properties indicate that a brain has structurally and functionally defined modules with short-ranged characteristics, where perturbations are localized, and then these local signals are propagated due to long-ranged interaction driven by rich club of key hubs. Moreover, weak interaction among these functional modules is essential for efficient information processing with minimum energy cost [4], and the dynamics and stabilization of the wiring/rewiring provide the basis for the adaptive nature of the brain network [6]. The modules are also arranged in a hierarchical manner with bunch of interacting key hubs exhibiting system level organization of the brain network, and through which a local information is distributed globally [5]. Further, the key hubs in the brain network are important for information integration and thus for performing complex cognitive functions, because these hubs are multifunctional, capable of long range information processing, and dynamically complex in nature [7]. However, still debatable open question is on how the energy management, distributions, and minimization happen among the interacting units (neurons or clusters of neurons) of a brain.

Study of a complex networks within the framework of Potts model was done in order to extract patterns and properties of the networks [13], and the formalism has been used as a powerful method for finding communities specially in hierarchical networks [14]. The simplified version of the Potts model, constant Potts model (CPM), in which Hamiltonian operator of the network is constructed from the evolved nodes and edges in the network, can be used to characterize various properties of the network. In this work, we used CPM in order to study properties of complex brain networks specially focusing on energy distributions in the network. Since brain networks follow hierarchical features [2, 8, 9], we apply the formalism to brain networks of *C. elegans*, cat, and macaque monkey [8], and studied how energy is distributed in the brain networks.

## Properties of Hamiltonian function in complex brain networks

The energy stored in a complex system represented by a complex network may be interpreted as the energy spent in the distribution and organization of interacting edges (wiring/rewiring of edges), with specified weights and directions, among the constituting nodes in the network. Hierarchical networks, which have the features of system level organization [10], involve the emergence of modules/communities which are also interacting in a certain fashion [11]. Each module/community has its own organization, consisting of cross-talking sub-modules/sub-communities in the next level of organization, which is less affected by sub-modules/sub-communities of other remaining modules/communities (Fig. 1). Similarly, the sub-modules/sub-communities will have their organization of sub-sub-modules/sub-sub-communities at the next further level, and so on. This system level organization of modules/communities at various levels amounts to the complexity in the network organization.

**Figure 1:**
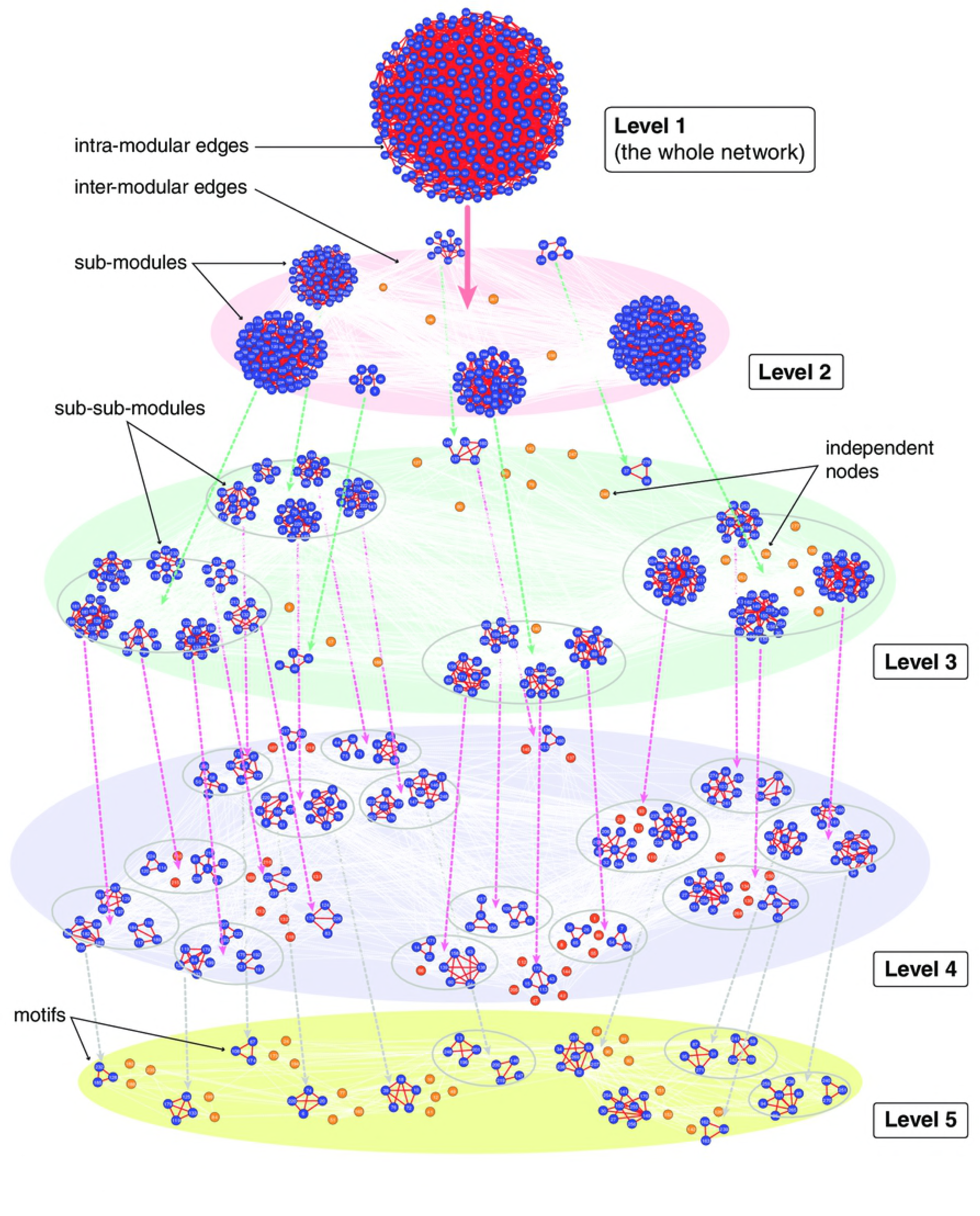
Hierarchically organized *C. elegans*’ neuronal network. Each layer (or level) constitutes a similar set of nodes and edges, but with different distributions describing various organizational hierarchies.

The modules/communities are relatively more densely connected groups of nodes, which could reflect different functions and cross-talk within the network. Here, we focus on the energy stored in a complex network at various levels of organization, and the way of using the stored energy in the network in organizing the network within the formalism of CPM [14]. Consider a network given by a graph, *G* = *G*(*E, V)*, with sets of constituting nodes *V* = {*k*}, *k* = 1, 2, …, *n*, and edges *E* = {*e*_*ij*_}, ∀*i, j* ∈*n*. The distribution of edges among the nodes is given by adjacency matrix *A*_*ij*_, ∀*i, j* ∈ *n*, such that *A*_*ij*_ = 1 if *i*th and *j*th nodes are connected, otherwise zero. The Potts model [12] can be used to analyze properties of complex network [13], and its simplified form, known as CPM, is used as a technique to detect communities in complex network [14]. For an undirected and unweighted network, the total number of edges in the network is *L* =∑_*ij*_ *A*_*ij*_, where *A* is *n* × *n* matrix. The Hamiltonian of this network within the CPM formalism [13–16] is given by

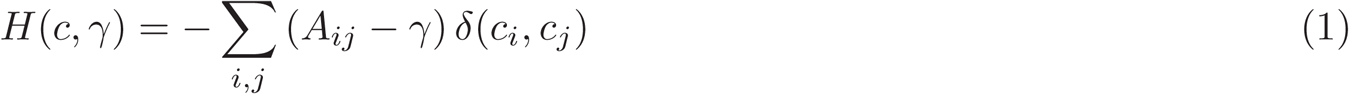

where *c*_i_ and *c*_j_ are *i*th and *j*th communities of the network, and *γ* is the resolution parameter of the network. Now rearranging equation (1), the Hamiltonian can be written as

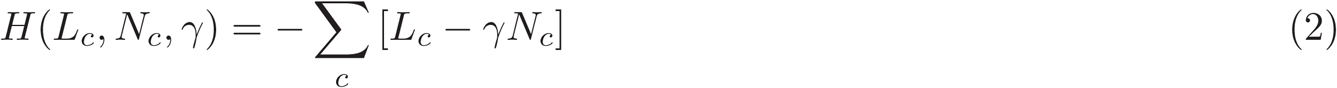

where *L*_*c*_ is the total number of internal edges in community *c*, and *N*_*c*_ = *n*_*c*_ × *n*_*c*_, with *n*_*c*_ is the total number of nodes in the community *c*. Complex natural networks are generally self-organized [17, 18], and have various levels of organization, with self-similar democratic constitution at each level of organization [19], down to the fundamental level where basic organizational units are motifs [20]. *C. elegans*’ neuronal network, consisting of *N* = 277 nodes, is one such type which have hierarchical organization of communities/sub-communities at various levels of organization (Fig. 1), and similar nature of topological organization was obtained in cat (*N* = 52) and macaque monkey (*N* = 71) brain networks also [8]. If such a network is defined by *G*(*L, N)*, then network at *level-2* is organized by a set of *m* communities defined by a set of sub-graphs 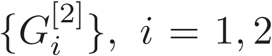, *i* = 1, 2, …, *m* constructed from *level-1*, i.e., the whole network. Since the networks at *level-1* and *level-2* represent the same set of nodes but with different network organization, we have 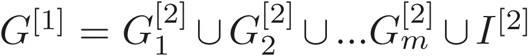, where *I*^[2]^ is the set of isolated nodes and their degrees. Hamiltonian at *level-1* is [1] given by, *H*^[1]^ = − (*L*^[1]^ − *γN* ^[1]^)) with 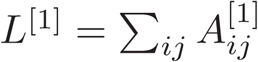 and *N* ^[1]^ = *n* × *n*. Now, the Hamiltonian of the network at *level-2* is given by 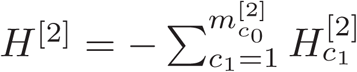, where *c*_1_th Hamiltonian at *level-2* is 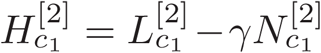 with 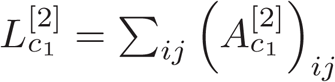, and 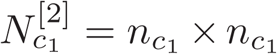 Similarly, organization of sub-communities in *level-3* is constructed from the respective communities in *level-2*. Therefore, Hamiltonian of the network at *level-3* can be derived as 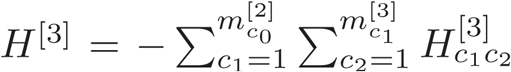, where Hamiltonian of *c*_2_ th sub-community at *level-3* derived from the *c*_1_ th community at *level-2* is 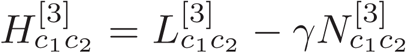 with 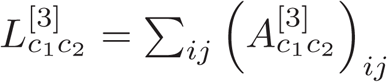, and 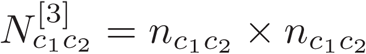. Hence, the Hamiltonian of the complete network *H*^[1]^ can be derived from the Hamiltonians at the lower levels (i.e., *level-2, level-3*,…,*level-U*) and is given by

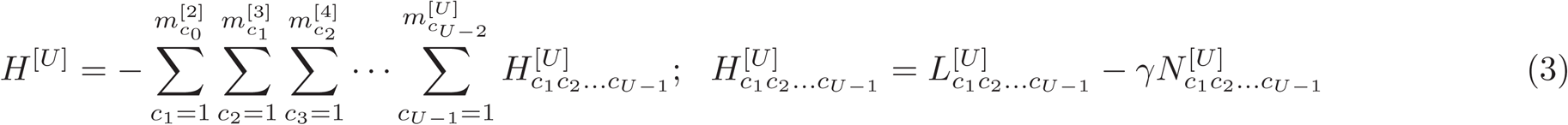

Here, 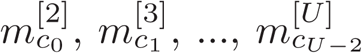 are numbers of communities at levels *level-2, level-3*,…,*level-U*, respectively. The sizes of the communities reduce as one goes from *level-1* to *level-U*, and this size reduction of communities with levels persists until the communities are reduced to motifs, which are building blocks of the network. Some communities have short histories of existence, and some have long histories. Nodes in communities having long histories generally have the tendency to regulate the network.

### Theorem 1.

In hierarchical networks having systems level organizations, the fluctuations in the number of nodes in going from one level of organization to the other (ΔN ^[k +1→k]^ = N ^[k]^ − N ^[k +1]^ satisfy the conditions:

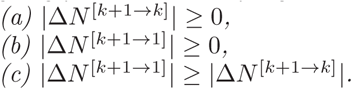

**Proof.** Equation (1) allows us to connect the size of the network at *level-1* to the size of the network at *level-2*, which is the sum of sizes of individual modules and isolated nodes at *level-2*, given by

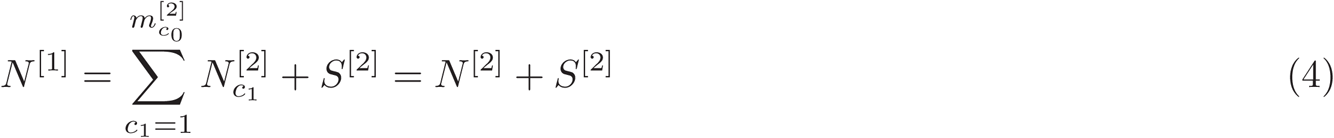

where 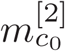 is the number of modules at *level-2*. 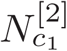 is the size of the *c*_1_ th module at *level-2*. *S*^[2]^ is the total number of isolated nodes at *level-2* of the network. The fluctuations in the number of nodes in retreating from *level-2* to *level-1* is given by |Δ*N* ^[2→1]^| = *N* ^[1]^ − *N* ^[2]^ ≥ 0 for *S*^[2]^ ≥ 0. The condition |Δ*N* ^[*k* +1→ *k*]^| = 0 satisfies only when *S*^[2]^ = 0, i.e., all the nodes at *level-1* are distributed to modules at *level-2*.

Proceeding in the same way, the number of nodes at *level-k* can be derived from the *level-(k+1)* expression using Equations (1) and (3):

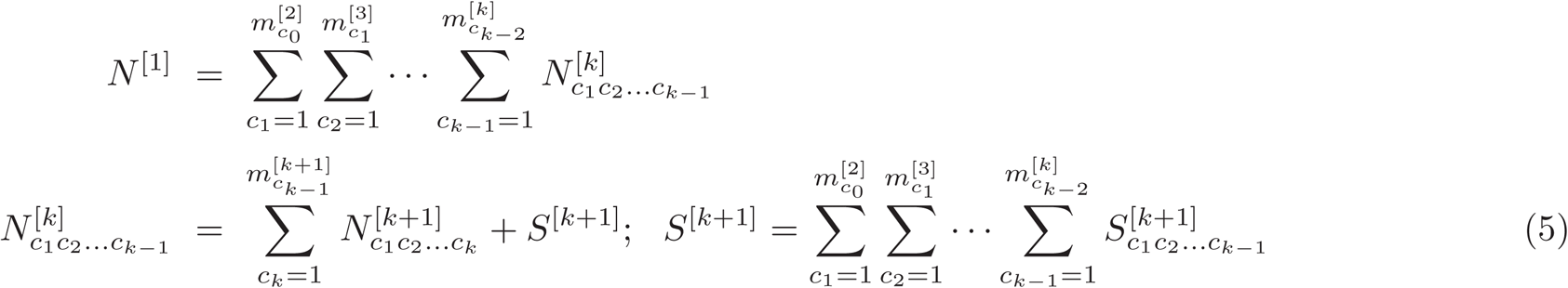

where 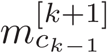 is the number of sub-modules at *level-(k+1)* constructed from *c*_*k*_ th module at *level-k*. 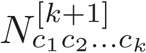 is the size of the *c*_*k*_ th sub-module at *level-(k+1)* derived from *c*_*k*-1_th module at *level-k*. If Δ*N* ^*k* +1→1^ is considered, then the sum over all nodes in modules/sub-modules and isolated nodes starting from *level-(k+1)* to *level-1* should be added. However, if Δ*N* ^*k* +1→ *k*^ is to be calculated then sum over all nodes in sub-modules and isolated nodes at *level-(k+1)* have to be done. This can be done by removing the sum indices *c*_1_ to *c*_*k*-1_ in Equation (5). The results are:

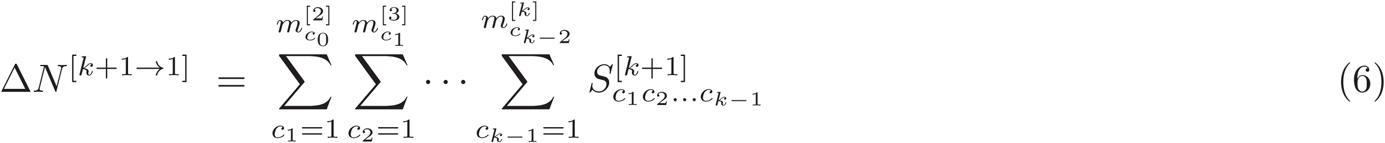

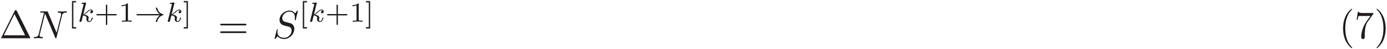

Then, it is trivial to prove that |Δ*N* ^[*k*+1→ *k*]^| = *N* ^[*k*]^ − *N* ^[*k* +1]^ = *S*^[*k* +1]^ ≥ 0 from Equation (7). Similarly, it can also be shown that |Δ*N* ^[*k* +1→1]^| ≥ 0 from Equation (6). It is also trivial from Equations (6) and (7) that |Δ*N* ^[*k* +1→1]^| ≥ |Δ*N* ^[*k* +1→1]^|. They are equal only when all the nodes in each level are totally distributed among the modules/sub-modules in the other level such that there are no isolated nodes.

### Theorem 2.

The variations in the number of edges as we move from one level of organization to the other (ΔL^[k +1→ k]^ = L^[k]^ − L^[k +1]^) because of wiring or rewiring among the modules satisfy the following conditions:

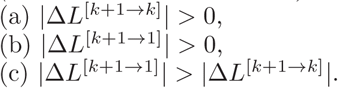

**Proof.** The total number of edges at *level-1* and *level-2* can be related by using Equation (1), as given below:

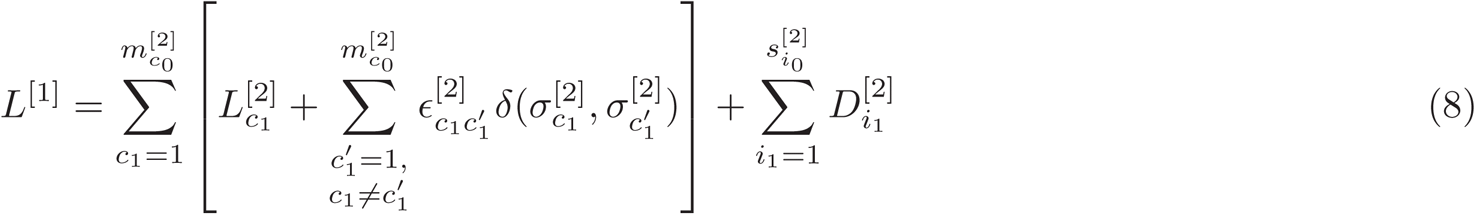

where 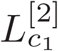 is the number of edges in the *c*_1_th module at *level-2*, constructed from the network at *level-1*. 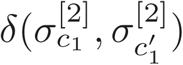 is delta function which is equal to one if modules 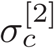 and 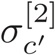 are connected, otherwise zero. 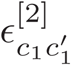 is the number of inter-edges between the connected *c*_1_th and 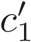 modules. 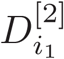 is the degree of *i*th isolated node at *level-2*, and 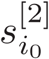 is the number of isolated nodes in the network at *level-2*. Since the first term is *L*^[2]^, we have

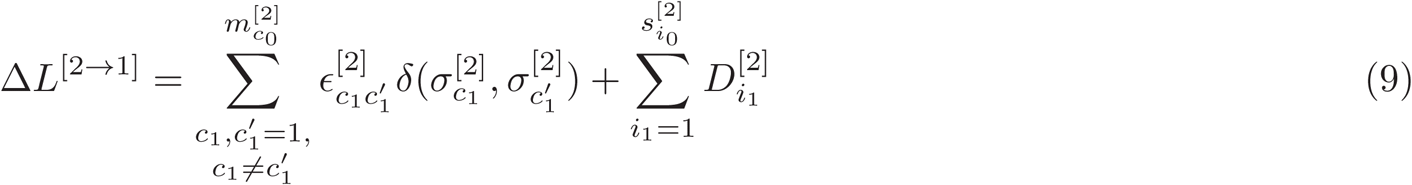

The second term could be zero if the nodes at *level-1* network are exactly distributed among the modules at *level-2*. The number of inter-edges among the connected sub-modules at *level-2* cannot be zero. Hence, |Δ*L*^[2→1]^| > 0.

Similarly, following the same process, we can generalize and arrive at the expressions for *L*^[1]^ constructed from the network organization at any *level-(k+1)*, and *L*^[*k*]^ constructed from neighboring level of organization *level-(k+1)*. The respective expressions are

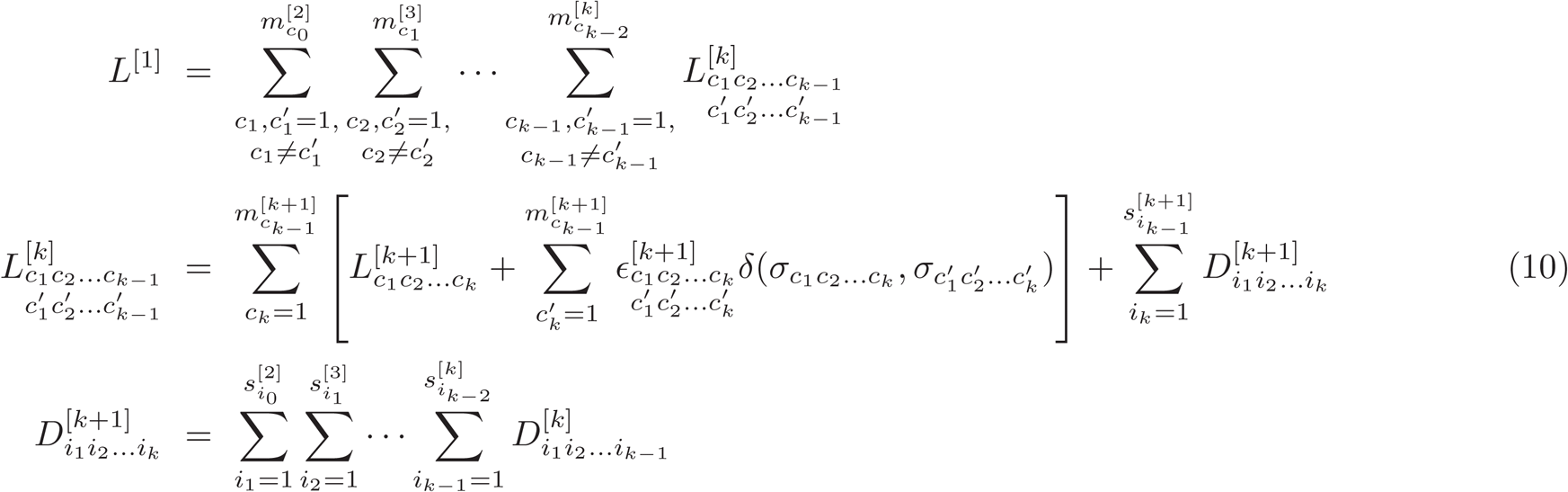

where 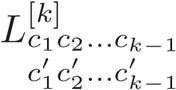 the number of edges of *c*_*k*–1_ th sub-sub-module at *level-k* constructed from *c* _*k*–2_ th sub-module at *level-(k-1)*, starting from *c*_1_th module at *level-2*. From Equation (10) we know that the total number of edges at *level-(k+1)* can be constructed from the edges at *level-k*. Then removing the running indices up to *c*_*k*-1_ in Equation (10), we get the fluctuations in edges when there is a change from *level-(k+1)* to *level-k* or vice versa. Then, the variations in edges due to changes from one level of organization to the other are given by

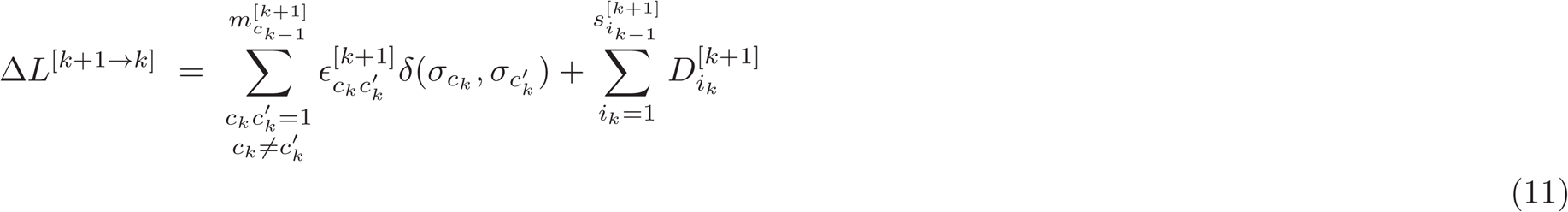

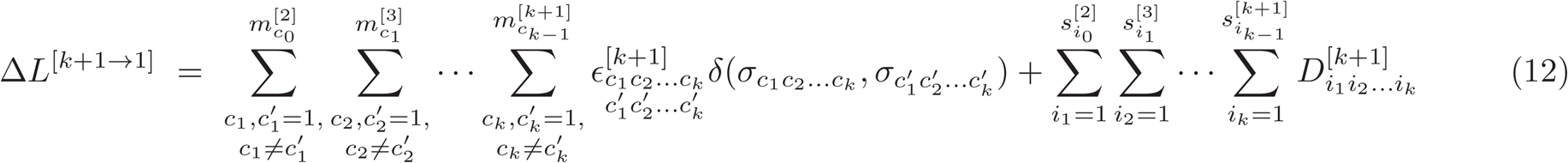

The second terms in Equations (11) and (12) are larger than zero if isolated nodes are involved when one crosses from one level of organization to another, otherwise the term equals to zero. However, the first terms in these equations are always greater than zero. This means the organization of a larger network at a particular level from a smaller network at another level needs significant number of edges for rewiring them to achieve the self-organization at that level. Hence, Δ*E*^[*k*+1→*k*]^ > 0 and Δ*E*^[*k*+1→1]^ > 0. Further, it is trivial that Δ*E*^[*k*+1→1]^ > Δ*E*^[*k*+1→*k*]^.

### Theorem 3.

The shift in the Hamiltonian of a complex network due to change of levels of organization satisfies |ΔH| > 0.

**Proof.** Equations (3), (6), (7), (11), and (12) allow us to write the expression for shift in Hamiltonian due to change in the levels of organization as

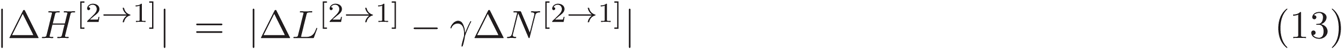

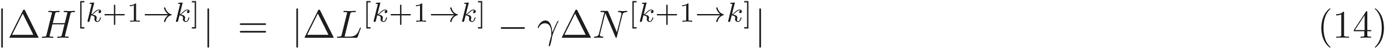

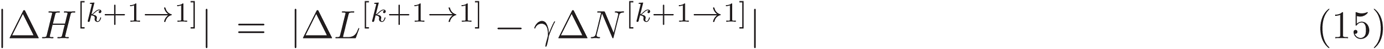

Since |Δ*L*^[2→1]^| > 0, |*L*^[*k*+1→*k*]^| > 0, |*L*^[*k*+1→1]^| > 0 |*N* ^[2→1]^| ≥ 0, |Δ*N* ^[*k*+1→*k*]^| ≥ 0, and |Δ*N* ^[*k*+1→1]^| ≥ 0, we can straightforwardly prove that |Δ*H*^[2→1]^| > 0. Further, it can also be proven that |Δ*H*^[*k*+1→*k*]^| > 0 and |Δ*H*^[*k*+1→1]^| > Also, there is a competition between the two terms Δ*L*^[*k*+1→*k*]^ and Δ*N* ^[*k*+1→*k*]^ magnified by resolution parameter *γ*. From Equations (6) and (12), we can further prove that |Δ*H*^[*k*+1→1]^| > |Δ*H*^[*k*+1→*k*]^|.

## Nature of Hamiltonian shift

Hamiltonian of a complex network is quantified by the contributions from the edges resulted from interactions among the nodes, and the mass of the network magnified by a resolution parameter *γ*. As this Hamiltonian function is related to the balances between the nodes and edges in a network, it is in fact related to the energy contained in the network. The organization of a network at various levels of organization have their own constitutions, which is most probably democratic (no emergence of central control mechanisms), and self-similar in nature. Since the organizational changes from on level of organization to another are connected, shifting one level of organization to another accounts to an energy cost for the organizational changes. To understand the behavior of Hamiltonian change due to wiring/rewiring in the network, we calculated the average Hamiltonian at each levels of organization (cf. Fig. 1) for the *C. elegans*, cat, and macaque monkey brain networks, and the same has been plotted as a function of level of organization (*s*) in Fig. 2 (upper three panels). Simulations for various values of resolution parameter *γ* are performed. There are two behaviors exhibited by these plots. The first behavior we can see from the curves is that in two phases they all follow power law behavior which is the signature of the self-organization in energy distributions. Hence, the behavior is given by ℍ (*s*) ∼ *s* ^Λ^, where 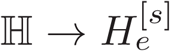 or 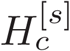 or 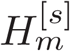 corresponds to power Λ → σ or δ or *ϵ*. A second observation from the plots is the variation in the sign and values of Λ as a function of *γ*: negative powers in Λ for large values of *γ* (Phase I), positive power values in Λ for very small values of *γ* (Phase III), and transition from negative values of Λ to positive values at moderate values of *γ* (Phase 2). These different phases driven by *γ* will have different properties in the distribution of energy. Hence, this resolution parameter *γ* could be one key parameter that reveals the energy distribution in brain networks.

**Figure 2:**
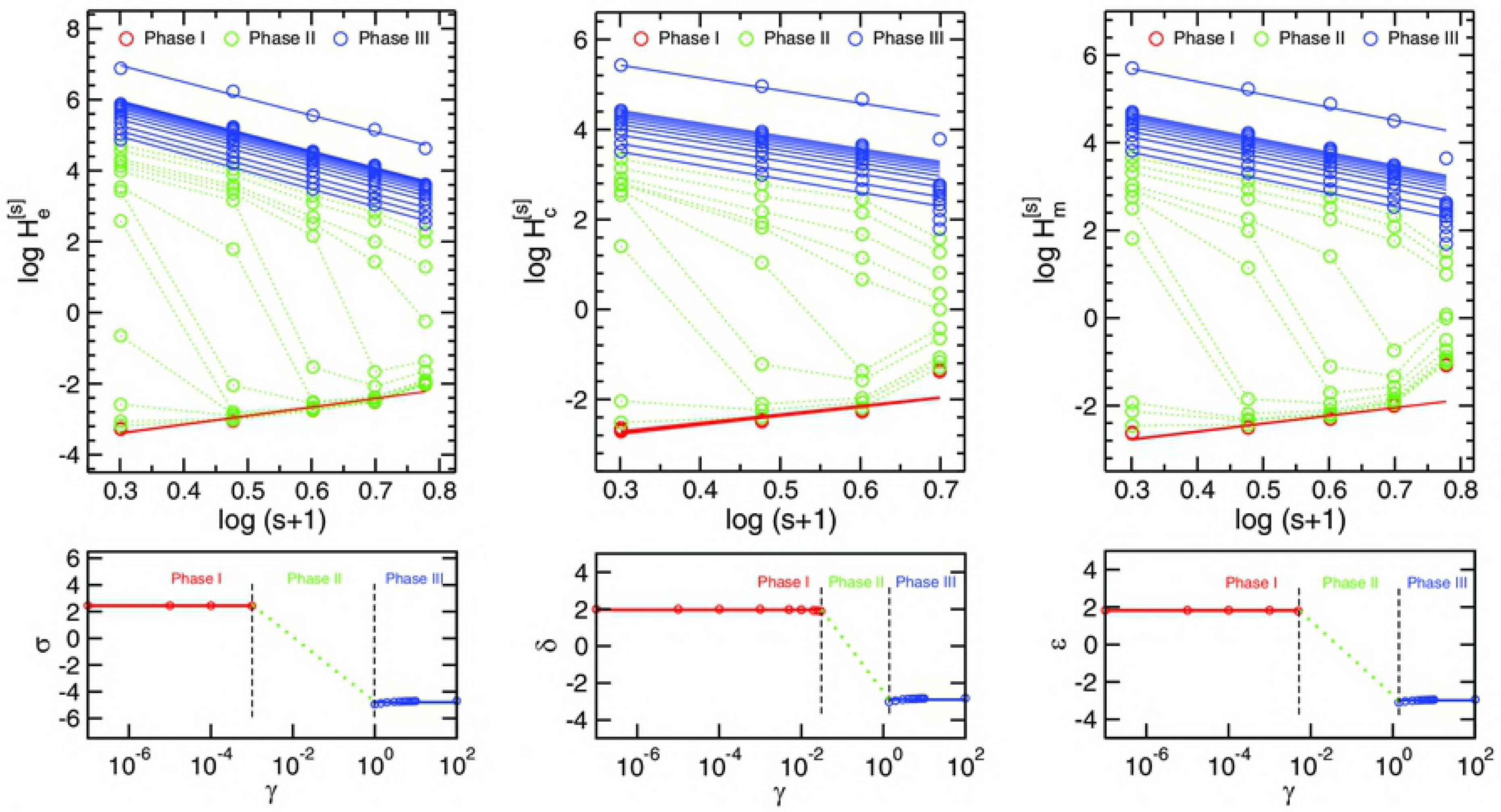
Hamiltonian spectrum of *C. elegans* (left), cat (middle), and macaque monkey (right) brain networks. Upper plots show the Hamiltonian function *H* as a function of levels of network organization *s* in log–log scale. Lower plots represent phase plots as a function of resolution parameter *γ*.

## Conclusion

One important quest in understanding variabilities in brain functionalities is how connectivities in complex brain networks get changed in random/non-random way with dynamic behavior [2]. Brain diseases which generally involve short and long range network dysfunctions could be due to the drastic change in the distributions of wiring/rewiring of these connectivities [4, 21]. These distributions of connectivities, in brain networks constructed from functional magnetic resonance imaging, diffusion tensor imaging, electroencephalogram, electrooculogram, etc., may be fundamentally related to energy distributions in brain. We have studied the energy distribution in the brain networks of three species [8] using simple but efficient CPM approach [12–14]. The edge distributions at various levels of organization are found to be different which are reflected in the energy distributions calculated via CPM, and also the Hamiltonian calculated as a function of levels of organization is found to follow power-law or fractal behavior which is a signature of self-organization red in the system. Moreover, the fractal nature of the Hamiltonian function is controlled by a key resolution parameter which balance the dominance of edges over nodes or vice versa. We also observe three different phases driven by this resolution parameter which may correspond to different brain states describing various situations/irregularities in the brain system. However, the existence of these brain states need to be verified experimentally.

Brain states with various phases exhibited are highly dynamic in nature which may depend on various factors. Since energy distributions among the the nodes in various communities and sub-communities could be one of the fundamental working principle in brain networks, studies in this perspective could give important insights of how brain system is organized.

## Acknowledgements

RKBS acknowledges CSIR, India, for providing financial support under sanction no. 25(0221)/13/EMR-II, MZM financially supported by Department of Health and Research, Ministry of Health and Family Welfare, Government of India under young scientist FTS No. 3146887. SSS is supported by SERB National Postdoctoral Fellowship (Grant No. PDF/2016/003188).

## Author Contributions

SKS, SSS, MZM, and RKB conceptualized the work. SKS, SSS, DH, and MZM did the simulation and preparation of the figures. SKS, SSS, DH, MZM, and RKB wrote the manuscript. All the authors read and approved the manuscript.

## Ethics approval and consent to participate

Not applicable

